# Benchmarking CUT&RUN analysis using motif enrichment

**DOI:** 10.64898/2026.09.09.749495

**Authors:** Luomeng Tan, Coby Viner, Xing Hao Li, Michael M. Wrana, Charles A. Ishak, Shu Yi Shen, Daniel D. De Carvalho, Sarah J. Hainer, Michael M. Hoffman

## Abstract

**Background:** Cleavage under targets and release using nuclease (CUT&RUN) maps the genome-wide locations of chromatin-associated proteins and provides an improved alternative to chromatin immunoprecipitation sequencing (ChIP-seq) for profiling sequence-specific transcription factor binding sites. Identifying these binding sites plays a critical role in understanding gene regulation, and transcription factors provide a useful setting for benchmarking because their well-defined sequence motifs serve as built-in controls for evaluating performance. Compared with ChIP-seq, CUT&RUN achieves higher resolution and lower background by avoiding cross-linking and bulk precipitation. Its distinct fragment length and cleavage characteristics, however, limit the direct transfer of existing computational tools, which primarily target ChIP-seq data. The performance of these tools on CUT&RUN can depend strongly on preprocessing choices. In this work, we investigate preprocessing strategies for transcription factor CUT&RUN, focusing on fragment length filtering and spike-in calibration. We aim to improve peak detection and provide practical guidance for analysis.

**Results:** We designed a benchmarking method to evaluate peak-calling procedures for CUT&RUN data and the effects of preprocessing approaches, including fragment length filtering and spike-in calibration. We benchmarked the two most widely used peak callers, MACS2 and SEACR, by assessing motif enrichment—the degree to which identified peaks contain the expected transcription factor binding motifs. Filtering for fragments with a length ≤120 bp generally improved target motif enrichment. Spike-in calibration using heterologous *Saccharomyces cerevisiae* DNA improved motif elucidation substantially for MACS2, with little benefit for SEACR. By contrast, using *Escherichia coli* DNA as a spike-in control often failed to produce valid results unless we could meticulously control *E. coli* contamination. MACS2 performed robustly across samples. SEACR performed especially well on clean, sparse-background datasets, but performed poorly on some datasets with denser background signal and often produced numerous apparent false positives. While MACS2 provided robust results under minor perturbations in fragment length filtering, SEACR exhibited greater sensitivity to such changes.

**Discussion:** Our benchmarking highlights how both peak caller choice and preprocessing strategy shape the analysis of transcription factor CUT&RUN data. By comparing the robustness and limitations of two widely used peak callers, we provide practical guidance on fragment length filtering, spike-in calibration, and tool selection. These findings help improve the processing and interpretation of CUT&RUN data, allowing researchers to more rapidly and reliably utilize this new technology. We expect that our work will guide more informed choices in CUT&RUN analysis and support the development of improved computational methodologies.

## Introduction

### Overview of CUT&RUN and biological motivation

Cleavage under targets and release using nuclease (CUT&RUN)^1^ maps the genome-wide locations of chromatin-associated proteins, including transcription factors, as well as histone modifications. By tethering a protein A-micrococcal nuclease (pA-MNase) fusion protein to antibodies bound to target proteins, CUT&RUN cleaves DNA near binding sites and releases short DNA fragments into solution.^1^ This targeted enzymatic cleavage eliminates the need for bulk ChIP and pull-down, which helps reduce background noise from non-specific DNA fragments and improves signal-to-noise ratio.^1^ Compared to chromatin immunoprecipitation sequencing (ChIP-seq),^2^ CUT&RUN typically achieves similar or greater efficiency with fewer cells and shorter processing time, making it a powerful tool for mapping protein–DNA interactions.^1^

Sequence-specific transcription factors regulate gene expression by binding to specific regulatory sequences in DNA.^3^ Identifying these binding sites plays a crucial role in determining the genes regulated by specific transcription factors and helps to understand the mechanisms of gene expression regulation.^4^ We therefore focused on transcription factors, for which well-defined binding motifs provide an internal benchmark for evaluating peak-calling performance.

For this purpose, CUT&RUN offers a more precise alternative to ChIP-seq. While ChIP-seq has long served as the standard assay for profiling protein–DNA interactions, its reliance on cross-linking and bulk precipitation introduces background noise and biases that limit resolution. By contrast, CUT&RUN directly cleaves near the antibody-bound sites, often yielding cleaner signal and more precise localization of transcription factor binding events.^1,5,6^

### CUT&RUN data characteristics and computational challenges

CUT&RUN aims to selectively cleave DNA at sites bound by targeted proteins, in our case, transcription factor binding sites. Variation in cleavage position relative to the binding site and surrounding chromatin structure leads to fragments of varying lengths. Fragment-size analysis of CUT&RUN data shows that cleavages around transcription factor binding sites predominantly yield small fragments (≤120 bp),^1^ likely representing direct transcription factor binding events. These fragments result from antibody-mediated recruitment of MNase, which cleaves DNA near the putatively bound transcription factor, typically flanking the binding site. Fragments with a length ≥150 bp generally correspond to nucleosome-protected regions.^1,7^ Nucleosomes shield adjacent DNA from cleavage and generate characteristic fragment sizes consistent with regular nucleosome spacing.

With few computational tools tailored for CUT&RUN data, many researchers adopt methods originally designed for ChIP-seq. Models underlying ChIP-seq analysis methods, however, often fail to account for the distinct biochemistry of CUT&RUN, which produces shorter fragments and lower background noise. As a result, the performance of these methods on CUT&RUN can depend strongly on data processing choices. As with any sequencing-based assay, CUT&RUN requires appropriate data processing to achieve reliable results.

Researchers have produced a small number of tools specifically tailored for CUT&RUN, such as CUT&RUNTools,^5,8^ a pipeline for quality assessment, analysis, and visualization, and the ssvQC^9^ R package for data quality assessment. Most prominently, the Sparse Enrichment Analysis for CUT&RUN (SEACR)^10^ software, designed alongside the laboratory assay, serves as a peak caller specifically for CUT&RUN and identifies regions with substantial mapped read enrichment. In a context with a growing availability of tools specific to CUT&RUN, we aim to provide a critical evaluation of strategies for data processing and analysis.

A key step in CUT&RUN data analyses involves peak calling—choosing an appropriate peak caller plays a pivotal role in obtaining accurate results. Among existing tools, Model-based Analysis of ChIP-Seq version 2 (MACS2),^11,12^ although designed for ChIP-seq data which typically exhibit higher background noise,^13^ remains widely used for CUT&RUN data. MACS2 empirically models the shifting size of ChIP tags to better localize precise binding sites and uses a local Poisson distribution of background signal to capture local bias.^13^ In ChIP-seq, these steps correct for random DNA shearing and substantial background noise, but in CUT&RUN, fragments arise from targeted MNase cleavage and already align closely with binding sites, producing much lower background noise. As a result, MACS2’s underlying assumptions may not fully capture the fragment and background properties of CUT&RUN data.

The typically low background signal in CUT&RUN data further increases the risk of false positives, as isolated background reads may appear as enriched regions.^10^ To address this challenge, SEACR provides a peak-calling approach tailored for datasets with low read depth and minimal background noise.^10^ SEACR uses a global empirical distribution of background signal to calibrate a simple threshold for peak calling, with the objective of minimizing false-positive peaks in comparison to other methods.^10^

### Data normalization and motif analysis in CUT&RUN

CUT&RUN, like all biological assays, contains sources of technical and biological variation that can obscure true signal, such as differences in sequencing depth and library complexity.^14,15^ Normalization ensures accurate data interpretation, but methods assuming equal total DNA yield across samples often fail because overall recovery varies between experiments.^16,17^ Spike-in calibration provides a solution by introducing heterologous DNA of known quantity whose yields remain unaffected by experimental conditions. The calibration process then entails normalization of spike-in read amounts across samples to identical levels, and the application of this same normalization coefficient to the experimental reads.^16^ By using spike-in calibration, researchers can normalize data to reveal changes or similarities hidden by experimental artefacts.^16^

To find transcription factor binding motifs from called peaks, researchers often perform down-stream motif analysis. Motif analysis plays a central role in assessing the biological relevance of identified peaks by testing for enrichment of known or de novo motifs near peak centres.^15^ The Centrality of Motifs (CentriMo)^18,19^ method performs motif enrichment analyses to identify motifs enriched in experimental data, leveraging the observation that the true binding sites in ChIP-seq experiments lie near the centres of the declared ChIP-seq peaks.^18^ Thus, the centrality of the target motifs in CentriMo results serves as an indicator of a successful experiment. CentriMo quantifies this enrichment using a metric called concentration—the total probability assigned to motif occurrences within a defined central region of the peak set.^18^ As increased concentration improves the accuracy of identifying the binding motif of the target transcription factor, we primarily utilize it across all of our benchmarking.

### Objectives of the study

Ultimately, achieving accurate results in genome-wide mapping assays requires careful consideration of multiple factors, including preprocessing strategies, accurate normalization methods, and selection of appropriate computational tools. These factors enable more accurate quantification of transcription factor binding and play a crucial role in ensuring reliable comparisons between samples. Here, we evaluate the performance of the two most commonly used peak callers for CUT&RUN data, MACS2 and SEACR, across a range of preprocessing strategies. We examine how fragment length filtering and spike-in calibration with *Saccharomyces cerevisiae* DNA and *Escherichia coli* DNA influence peak detection and the interpretability of data.

## Results

### Overview

We designed a benchmarking method (Figure 1; see Methods) to evaluate peak callers and the influence of fragment length filtering and spike-in controls on transcription factor binding elucidation. We collected 12 CUT&RUN samples from six laboratories (Table 1). We analyzed each sample with a corresponding immunoglobulin G (IgG) control sample as a negative control to assess background or non-specific binding. We ran the benchmarking workflow eight times on each sample, based on all conditional combinations (Figure 2). We assessed each combination’s performance based on the centrality of the target motif within peak sets determined by CentriMo concentration.

**Figure 1.**
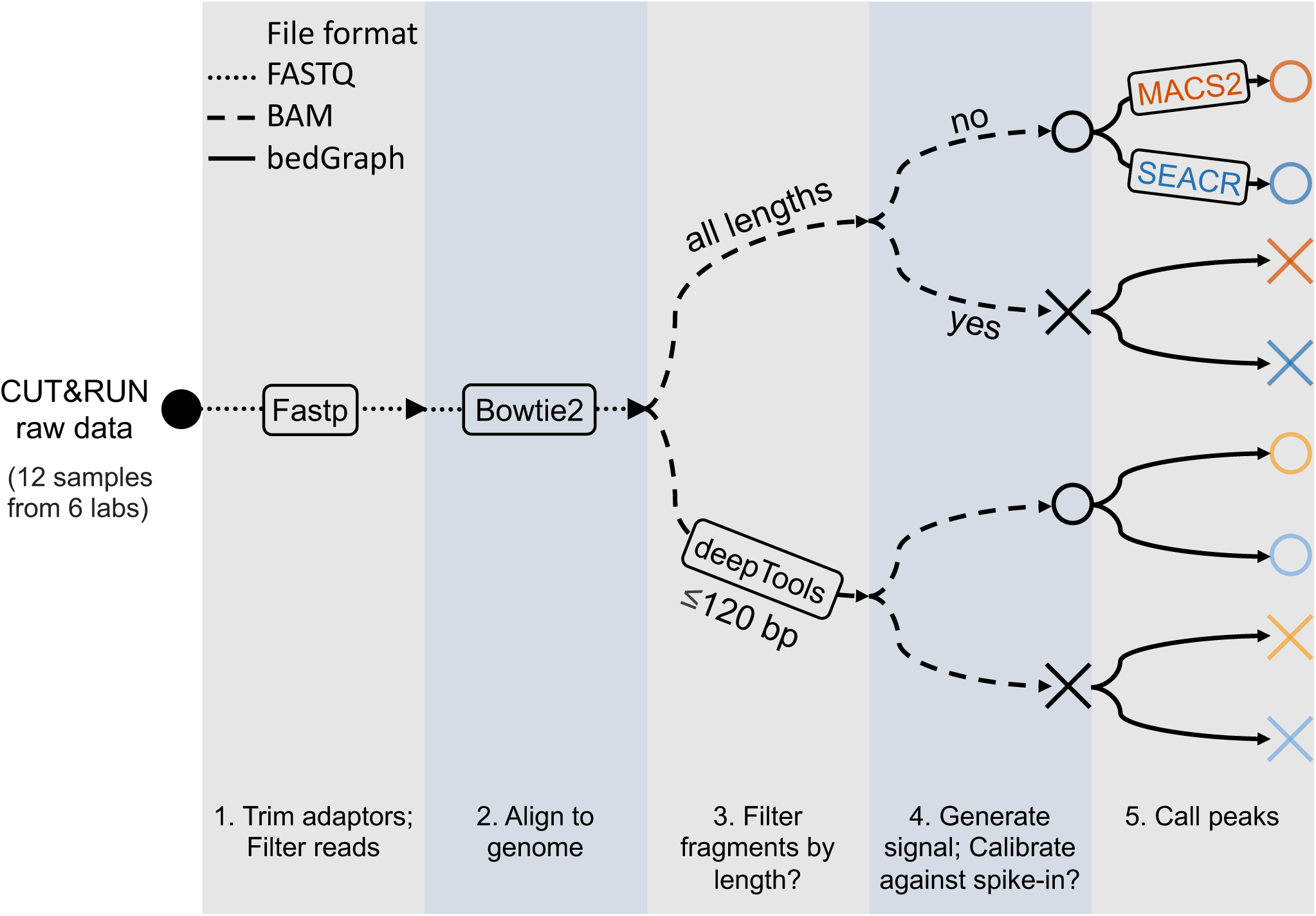
Overview of the benchmarking method. Our pipeline accepts paired-end CUT&RUN data as input, performs peak calling, and produces a set of peaks that denote putative transcription factor binding sites. Color hue: peak caller (orange: MACS2, blue: SEACR). Light colors: fragment length filtering ≤120 bp; dark colors: no fragment length filtering. Crosses: the use of spike-in calibration; circles: no calibration.

**Table 1.** Datasets used for benchmarking. Each dataset lists the sample (the target transcription factor and sample number (#), if applicable), Gene Expression Omnibus (GEO) accession, reference genome assembly used for alignment, and cell type. ESCs: embryonic stem cells.

| Lab | Sample | GEO<sup>20</sup> accession | Assembly | Cell type |
| --- | --- | --- | --- | --- |
| De Carvalho | C/EBPβ#1 | GSM9863663 | GRCh38/hg38<sup>21</sup> | K562 cells<sup>22</sup> |
| | C/EBPβ#2 | GSM9863664 | | |
|  | ZNF143 | GSM9863665 |  |  |
| Hainer | CTCF | GSM3022415<sup>23</sup> | GRCm38/mm10<sup>24</sup> | E14 mouse ESCs |
|  | SOX2 | GSM3022432<sup>23</sup> |  |  |
| Henikoff | FOXA2<sup>a</sup> | GSM3609741<sup>25</sup> | GRCh37/hg19<sup>26</sup> | Definitive endoderm<sup>b</sup> |
|  | SOX2<sup>a</sup> | GSM3609746<sup>25</sup> |  | H1 human ESCs<sup>27</sup> |
|  | CTCF | GSM2803196<sup>28</sup> |  | K562 cells |
| Roberts | ESRRB | GSM5703789<sup>29</sup> | GRCm38/mm10 | Mouse ESCs |
| Vakoc | NFYB | GSM7103770<sup>30</sup> | GRCh38/hg38 | RH4 cells<sup>31</sup> |
|  | NFYC | GSM7103771<sup>30</sup> |  |  |
| Wysocka | ALX4 | GSM7213748<sup>32</sup> | GRCh38/hg38 | H9 human ESCs<sup>27</sup> |

<sup>a</sup> Dataset called “gold standard” in the SEACR study<sup>10</sup>
<sup>b</sup> Derived from H1 human ESCs

### Fragment length filtering improves transcription factor binding elucidation

To examine whether filtering based on fragment length improves transcription factor binding signal detection, we filtered the reads from the C/EBP*β*#1 sample using a 120 bp cut-off, as recommended in the CUT&RUN protocol.^1,7^ We compared motif enrichment in data filtered by read length ≤120 bp versus unfiltered data. The CentriMo results show significant central enrichment (concentration: 0.0823) of the target C/EBP*β* motif in filtered fragments (Figure 3b), while unfiltered fragments show no significant central enrichment (concentration: 0.0435; Figure 3a). A flat distribution in the unfiltered data suggests that target motif occurrences follow a uniform distribution rather than showing central enrichment. We observe similar behaviours for FOXA2 (Figure 4a–b). The curve shapes for SEACR peaks before and after filtering show similarity, but the probability at the central position (0) increases from 0.0798 for unfiltered fragments (Figure 4c) to 0.0943 for filtered fragments (Figure 4d). In summary, fragment length filtering enhanced peak detection by both MACS2 and SEACR, yielding stronger central motif enrichment.

We filtered all samples for fragments with lengths ≤120 bp. Fragment length filtering increased the concentration for all samples using MACS2 and for 8/12 samples for SEACR (Figure 2). For the 4 samples where filtering did not increase concentration, two of them simply had 0 concentration using SEACR (C/EBP*β*#1 and ZNF143). This likely resulted from SEACR’s design and optimization for sparse-background data, whereas these two samples showed relatively dense background signal. For the other two samples, filtering resulted in lower concentration values (0.2589 → 0.2278 for NFYB; 0.2969 → 0.2571 for NFYC). Fragment length filtering can have complex impacts on binding site elucidation and central enrichment (see Discussion).

We also investigated changes in peak number after fragment length filtering. We observe a negative correlation (Pearson *r* = −0.27; *p* < 0.05) between peak number and concentration, with peak sets containing a target motif concentration below 0.06 tending to show large numbers of peaks (Figure 2, top). This pattern suggests that many of these peaks likely represent false positives. Although both peak callers occasionally generated large peak sets at low target motif concentrations, this occurred more prominently for MACS2 without fragment length filtering. This pattern likely arises because MACS2 uses a background to locally distinguish signal from noise, while CUT&RUN’s sparse background can cause MACS2 to spuriously call some regions with weak signals as peaks.

**Figure 2.**
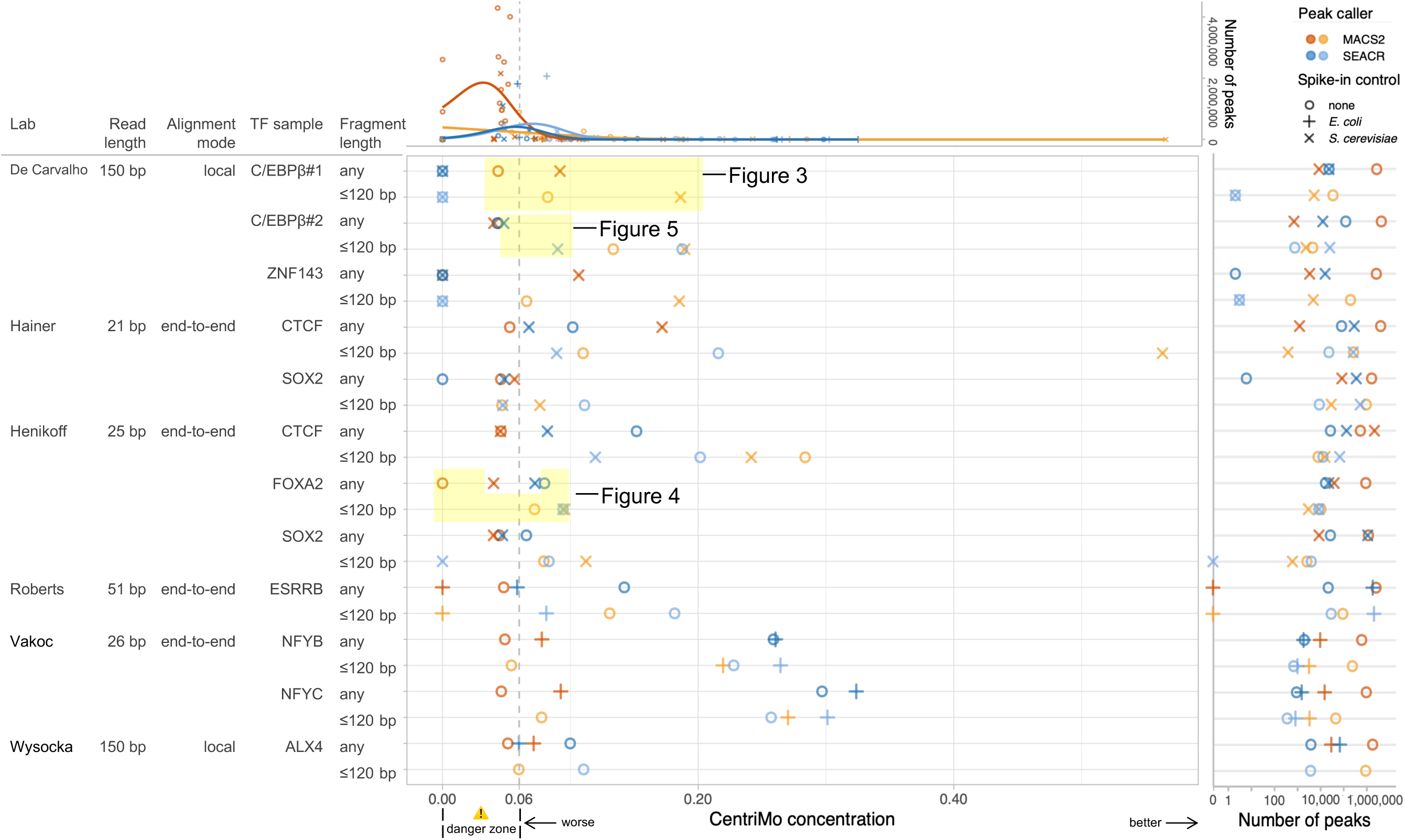
Summary plot: known transcription factor motifs elucidated by different methods. CentriMo concentration refers to the total probability assigned to target motif occurrences within the central 20 bp region. Pairs of rows denote results for each sample (top: no fragment length filtering; bottom: retaining only fragments ≤120 bp). Each row includes concentration values for all four combinations of peak caller and spike-in calibration method. The ALX4 sample with fragments ≤120 bp includes only two combinations, as insufficient spike-in reads prevented spike-in calibration. Colour hue: peak caller (orange: MACS2, blue: SEACR). Light colors: fragment length filtering ≤120 bp; dark colors: no fragment length filtering. Shape: no spike-in control (circle); *Escherichia coli* spike-in control (plus); *Saccharomyces cerevisiae* spike-in control (cross). Each sample used only one spike-in organism. *(Left)* Sample metadata, sequencing, and alignment characteristics. We aligned >80 bp samples using local mode and aligned others end-to-end. *(Center, top)* Number of peaks against CentriMo concentration. The curves depict a generalized linear model and highlight the trend for the two peak callers. *(Center, bottom)* CentriMo concentration. We used an empirically calculated concentration threshold of 0.06 as indicative of significant central enrichment. We classified any sample with a CentriMo concentration of 0.00–0.06 as “in the danger zone”: these samples often exhibited non-specific binding to target motifs, making them unsuitable for drawing any particular conclusions. Highlighted peak sets have site probability plots elsewhere in results for C/EBP*β* (Figures 3, 5) and FOXA2 (Figure 4). *(Right)* Number of peaks called.

Fragment length filtering helps remove these regions with weak or spurious signals by excluding longer fragments less likely to contain true binding sites. We observed a large reduction in the number of peaks for MACS2 after applying fragment length filtering across all samples, with a median reduction of 1 044 886 peaks, accompanied by a median increase in concentration of 0.0481. SEACR peaks showed a similar reduction in 9/12 samples, with a median reduction of 4562 peaks and a median increase in concentration of 0.0161. These results underscore the importance of fragment length filtering for accurate data interpretation and optimal motif elucidation.

In addition, fragment length filtering eliminated the bimodal distribution observed in the CentriMo plot (Figure 5)—which typically signifies an anomaly—and increased the concentration from 0.0479 to 0.0901. Bimodal peaks may arise when MNase cleaves two proximal binding sites, possibly due to non-specific binding or background cleavage.

To assess whether alternative fragment length thresholds might outperform the commonly used 120 bp cut-off, we tested a range of thresholds from 90 bp to 140 bp (Figure 6). To manage computational costs, we selected two representative datasets: the C/EBP*β*#1 (where the 120 bp cut-off resulted in a concentration of 0 using SEACR) and FOXA2 (a benchmark sample from the SEACR paper^10^). While the 120 bp cut-off generally improved concentration, it did not always yield optimal results. Notably, using a slightly different threshold restored target motif enrichment for C/EBP*β* with SEACR (a 121 bp cut-off resulted in a concentration of 0.1991), indicating that the poor result using a 120 bp cut-off arose from threshold sensitivity rather than sample quality. We further discuss this threshold sensitivity in a later section.

Smaller cut-offs tend to increase concentration further, particularly for MACS2 (Figure 6). Reducing fragment length beyond a certain point, however, may lead to a drop in concentration and limit biological relevance, as it may filter out fragments associated with genuine transcription factor binding. Given our current understanding of the underlying biochemistry and the widespread use of a 120 bp cut-off, we selected this threshold as a practical default for our benchmarking analyses. Moreover, while an alternative threshold rescued SEACR performance in this specific case (C/EBP*β*#1 sample), the erratic trend observed in SEACR’s concentration curve suggests that this variability reflects a limitation of the tool itself, rather than a need for sample-specific threshold optimization. The 120 bp threshold remains broadly effective and interpretable.

**Figure 3.**
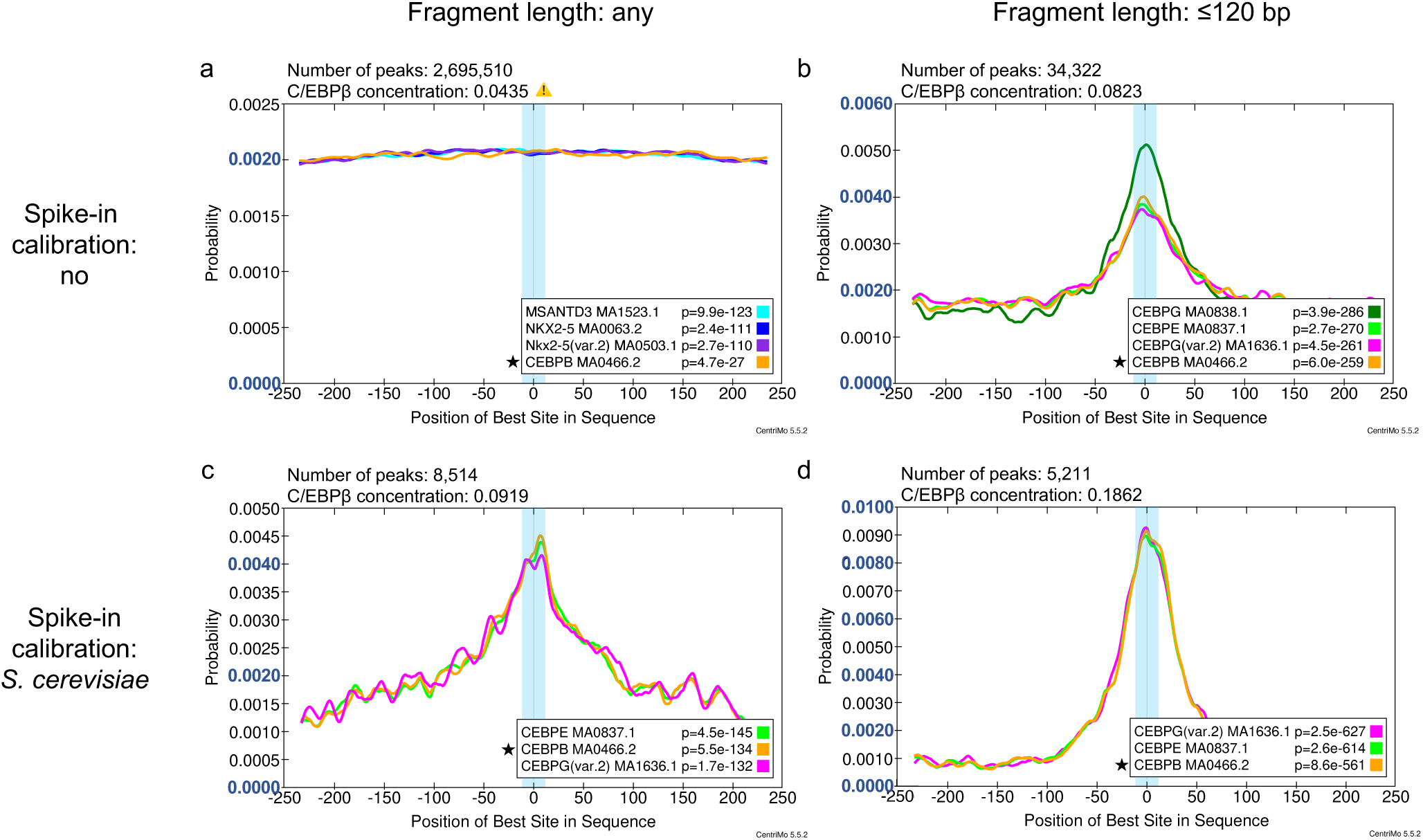
Filtering for fragments with lengths ≤ 120 bp and applying spike-in calibration led to greater central enrichment of the C/EBP*β* motif. All plots depict motif probability curves for the top three motifs, ranked by p-value, and the target motif (if not within the top three). The plots show the probability of the best match to a given motif against sequence position. Orange and a star (★) prefix mark the target motif, C/EBP*β*. Data come from the C/EBP*β*#1 sample. We only presented results for MACS2 peaks and omitted SEACR results, since for SEACR, the target C/EBP*β* motif had no enrichment. The y-axis scale differs across the plots. We use blue colouring to demarcate every 0.0020 increase in probability. 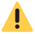: concentration in the danger zone. **(a)** Before spike-in calibration for all fragments. **(b)** Before spike-in calibration for fragments with lengths ≤120 bp. **(c)** After spike-in calibration using *Saccharomyces cerevisiae* for all fragments. **(d)** After spike-in calibration for fragments with lengths ≤120 bp.

**Figure 4.**
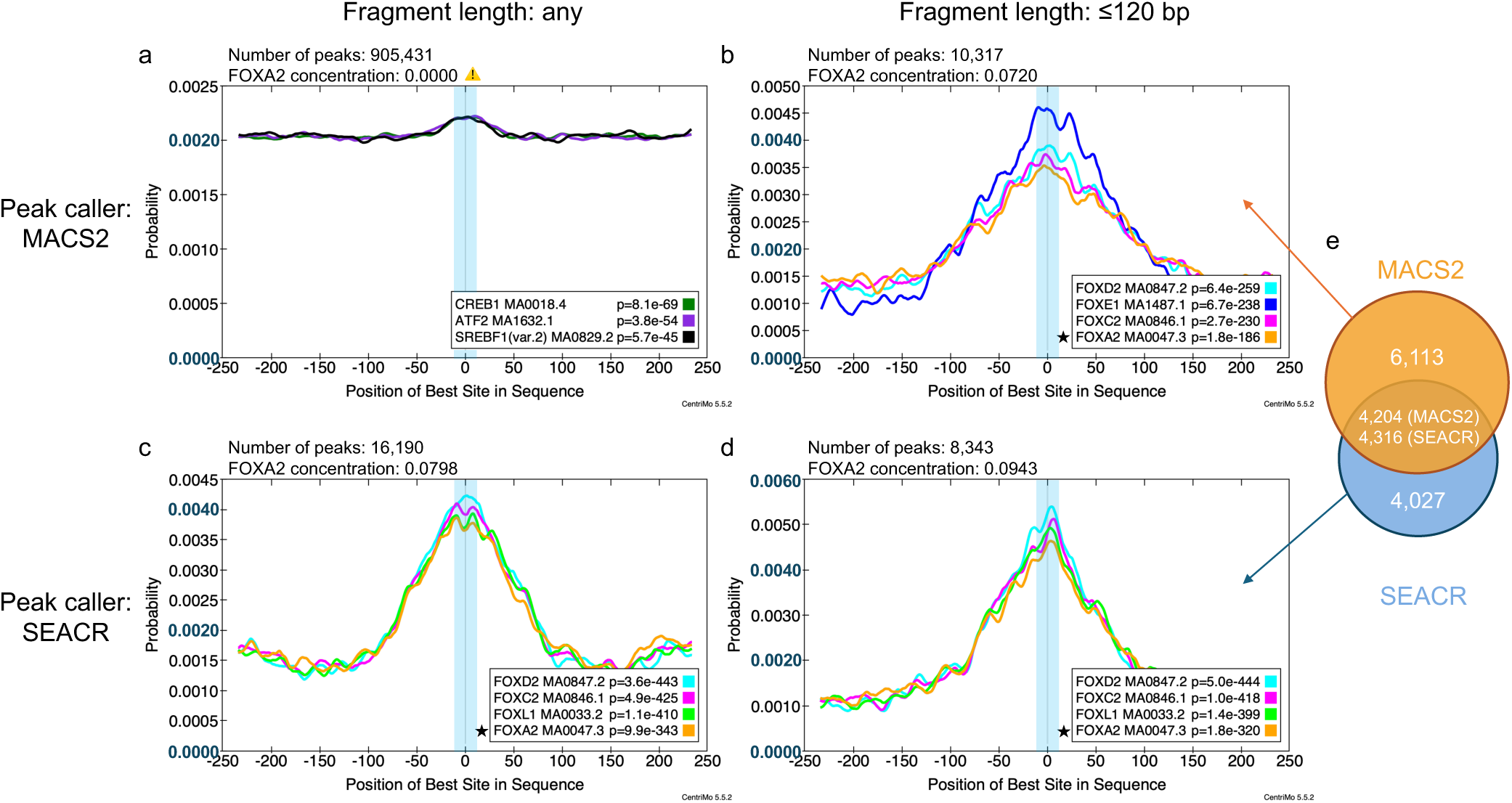
Fragment length filtering led to greater central enrichment of the FOXA2 motif for both MACS2 and SEACR peaks. All plots depict motif probability curves for the top three motifs, ranked by p-value, and the target motif (if not within the top three). The plots show the probability of the best match to a given motif against sequence position. Orange and a star (★) prefix mark the target motif, FOXA2. We did not apply spike-in calibration for these results. The y-axis scale differs across the plots. We use blue colouring to demarcate every 0.0020 increase in probability. 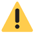 : concentration in the danger zone. **(a)** MACS2 peaks for all fragments. **(b)** MACS2 peaks for fragments with lengths ≤120 bp. **(c)** SEACR peaks for all fragments. **(d)** SEACR peaks for fragments with lengths ≤120 bp. **(e)** Overlap summary of MACS2 and SEACR peaks using fragments ≤120 bp. We considered MACS2 and SEACR peaks as common if either showed at least a 50 % overlap in included bases with the other. The non-overlapping regions indicate unique peaks, whereas the shared region reports the numbers of peaks classified as common for each caller. Because a peak from one caller could overlap multiple peaks from the other caller, the number of common peaks differed between MACS2 and SEACR.

**Figure 5.**
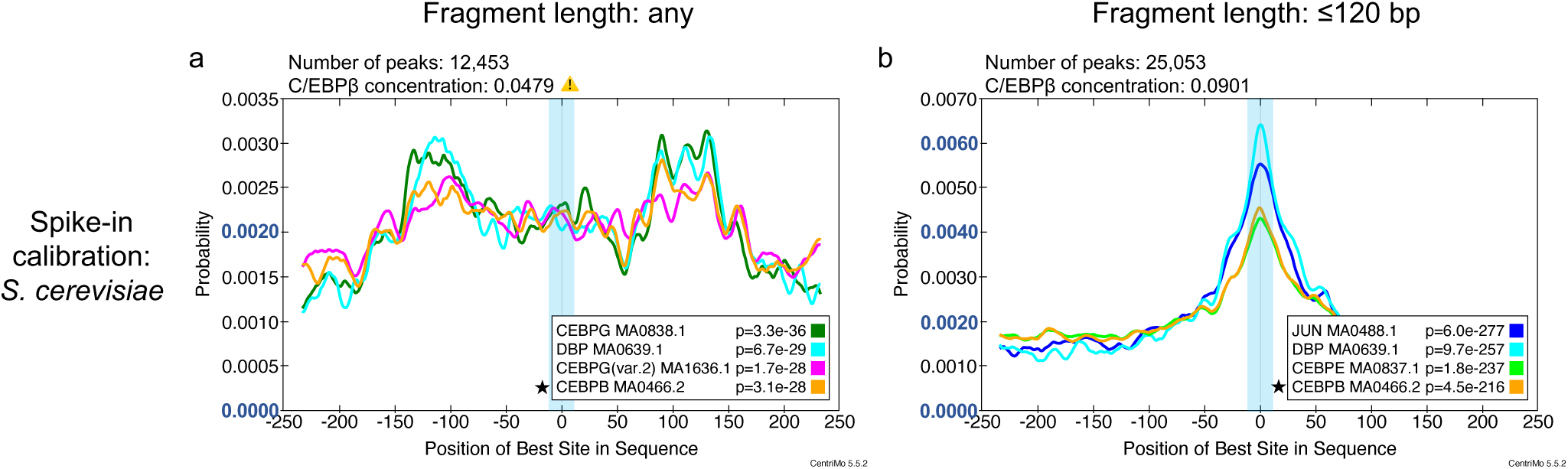
Filtering for fragments with lengths ≤120 bp removes the bimodal positional enrichment pattern. Data come from the second C/EBP*β* sample. All plots depict motif probability curves for the top three motifs, ranked by p-value, and the target motif (if not within the top three). The plots show the probability of the best match to a given motif against sequence position. Orange and a star (★) prefix mark the target motif, C/EBP*β*. The y-axis scale differs across the plots. We use blue colouring to demarcate every 0.0020 increase in probability. 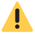 : concentration in the danger zone. **(a)** All fragments. **(b)** Fragments with lengths ≤120 bp.

**Figure 6.**
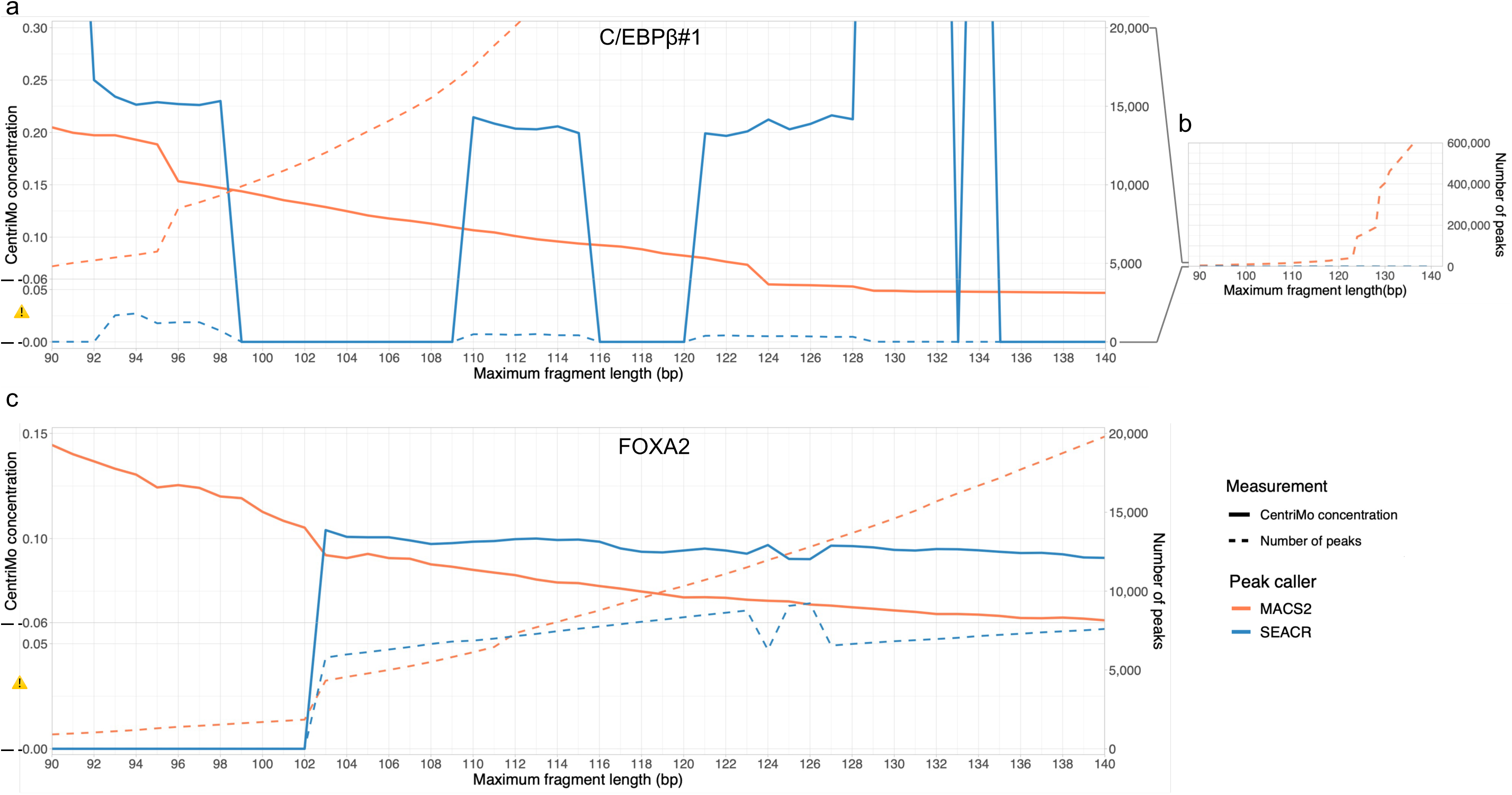
CentriMo concentration and peak number using different cut-offs for fragment length filtering. The concentration of target motifs and ensuing peak numbers when filtering for fragment length with cut-offs ranging from 90 bp to 140 bp for MACS2 (orange) and SEACR (blue). An empirically calculated concentration threshold of 0.06 represents significant central enrichment. We did not employ any spike-in calibration for these analyses. 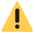 : concentration in the danger zone. **(a)** C/EBP*β*#1, showing CentriMo concentration (solid lines, left y-axis) and peak numbers (dashed lines, right y-axis), with the peak-number y-axis limited to 20 000. **(b)** C/EBP*β*#1, showing the full range of peak numbers. **(c)** FOXA2, showing CentriMo concentration (solid lines, left y-axis) and peak numbers (dashed lines, right y-axis).

### Spike-in calibration boosts peak calling specificity

We evaluated the effect of spike-in calibration using control DNA from *S. cerevisiae* and *E. coli*. Spike-in calibration revealed the central enrichment of target motifs for C/EBP*β*#1, with concentration increasing from 0.0435 without spike-in calibration to 0.0919 with spike-in calibration (Figure 3c). When combined with fragment length filtering, concentration further increased to 0.1862 (Figure 3d). Spike-in calibration, especially with *S. cerevisiae* spike-ins, however, only increased the concentration when calling peaks with MACS2—concentration increased for MACS2 for 9/12 samples, but only for 2/12 SEACR samples (Figure 7).

Our results suggest that using *E. coli* spike-ins presents challenges unless we meticulously control *E. coli* contamination, which often proves impractical. Both NFYB and NFYC samples showed an unexpectedly high number of *E. coli* spike-in mapped reads, with 5 316 628 reads in NFYB and 6 242 768 reads in NFYC after fragment length filtering. Similarly, the ESRRB sample contained a large number of putative *E. coli* reads (124 198 reads after fragment length filtering), differing substantially from the IgG control sample generated in parallel (108 reads after fragment length filtering). Given the difficulty in distinguishing between some possible acceptable or usual level of *E. coli*, versus substantive contamination, or potentially spiked-in bacterial DNA, we cannot recommend *E. coli* in spike-in calibration. Its use tends to yield inaccurate and unpredictable results. The arbitrary scale value used in computing spike-in factors greatly influenced MACS2’s performance. We typically use 10 000, as initially suggested.^28^ This scale, however, did not always produce optimal results, making it necessary to carefully select an appropriate arbitrary scale for each sample. We depict the results for the C/EBP*β*#2 sample, using various arbitrary scales as an example (Table 2). Here, we calculated the spike-in factors using spike-in read counts after restricting fragments to lengths ≤120 bp. Table 2 therefore does not represent the conventional arbitrary-scaling approach described in Methods, which uses unfiltered spike-in read counts. Instead, it illustrates how the choice of arbitrary scale affects performance within the filtered workflow.

When the arbitrary scale greatly exceeded the spike-in read count, the resulting spike-in factor became large and produced numerous non-specific peaks. As the arbitrary scale increased from 50 to 10 000, the number of peaks rose from 287 to 731 704, while the concentration dropped from 0.1976 to 0.0473. This sharp increase in peak number, together with the drop in concentration, suggests that many of the additional peaks likely reflected false positives. By contrast, specificity improves when the arbitrary scale closely matches spike-in counts—for example, a scale of 100 yields 1040 peaks with a concentration of 0.2167.

**Figure 7.**
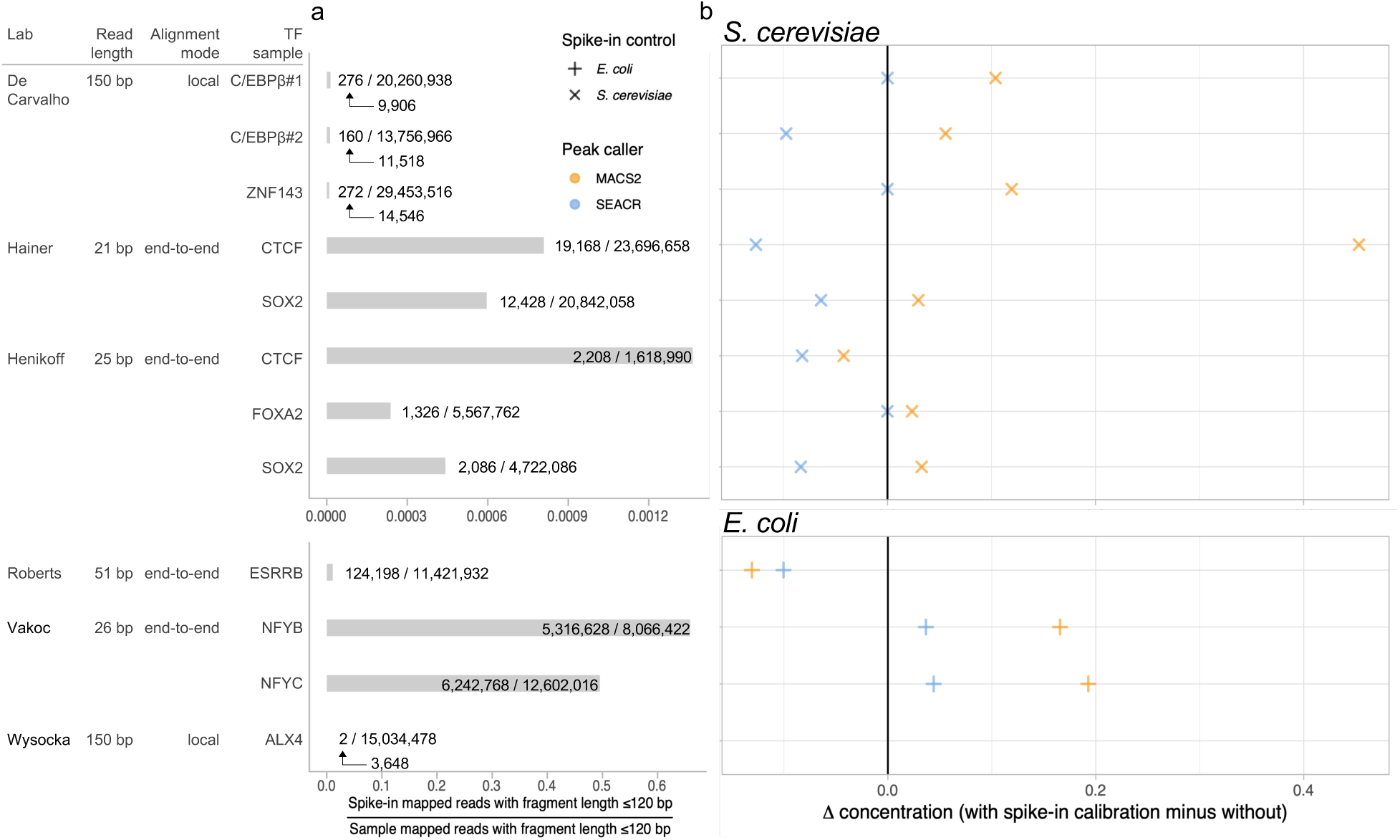
Effects of spike-in calibration on different peak callers. Each row pertains to samples using *Saccharomyces cerevisiae* (top) or *Escherichia coli* spike-ins (bottom). We filtered for fragment lengths ≤120 bp. Colour hue: peak caller (orange: MACS2, blue: SEACR). Shape: no spike-in control (circle), *Escherichia coli* spike-in control (plus), *Saccharomyces cerevisiae* spike-in control (cross). **(a)** Number of reads mapped to spike-in as a fraction of reads mapped to the sample. For the samples with the fewest reads mapped to spike-in, we also indicate the number of reads mapped to spike-in before fragment length filtering, with arrows connecting these values to the corresponding filtered counts. **(b)** Difference between the concentrations obtained with spike-in calibration and those without. Concentration for ALX4 not shown due to insufficient spike-in reads for calibration.

**Table 2.** Comparison of the number of peaks in the C/EBP*β*#2 dataset versus target motif concentrations, between MACS2 and SEACR, across different spike-in factors. To calculate the spike-in factor, we divided the arbitrary scale by the number of spike-in reads. The C/EBP*β*#2 sample contained 160 spike-in mapped reads, and the IgG control sample contained 276. We filtered for fragments lengths ≤120 bp. The bold number highlights the highest concentration observed for MACS2.

| Scale<sup>a</sup> | Spike-in factor<sup>b</sup> |  | MACS2 |  | SEACR |  |
| --- | --- | --- | --- | --- | --- | --- |
| | C/EBPβ | IgG | # peaks | conc<sup>c</sup> | # peaks | conc |
| 10 | 0.06 | 0.04 | 7 | NA<sup>d</sup> | 25 053 | 0.0901 |
| 50 | 0.32 | 0.18 | 287 | 0.1976 | 25 053 | 0.0901 |
| 100 | 0.62 | 0.36 | 1040 | **0.2167** | 25 053 | 0.0901 |
| 200 | 1.25 | 0.72 | 9088 | 0.1299 | 25 053 | 0.0901 |
| 500 | 3.12 | 1.82 | 7968 | 0.1358 | 25 053 | 0.0901 |
| 1000 | 6.25 | 3.62 | 99 800 | 0.0588 | 25 053 | 0.0901 |
| 2500 | 15.62 | 9.06 | 103 381 | 0.0587 | 25 053 | 0.0901 |
| 5000 | 31.25 | 18.12 | 731 267 | 0.0471 | 25 053 | 0.0901 |
| 10 000 | 62.50 | 36.24 | 731 704 | 0.0473 | 25 053 | 0.0901 |

<sup>a</sup> Arbitrary scaling coefficient
<sup>b</sup> Scale inconsistency arises solely from rounding.
<sup>c</sup> CentriMo concentration
<sup>d</sup> No significant central enrichment for the target motif.

Selection of an arbitrary scale lacks a clear standard. This makes it difficult to determine the appropriate value and can introduce inconsistency between samples. To address this, we undertook an empirical scaling factor calibration method: we performed spike-in calibration by scaling down the larger sample to match the smaller one, rather than relying on an arbitrary scale (Methods). We also applied the same fragment length filtering to spike-in reads as employed for target reads.

Compared to using an arbitrary scale of 10 000 and unfiltered spike-in read counts for all conditions, this new spike-in calibration method improves results for MACS2 for 7/12 samples that initially showed poor concentration (≤0.06) for the target motif (Table S1). Applying this method also improves the SEACR results for one sample from an initial concentration of 0 to 0.0901. This improvement likely resulted from the use of filtered spike-in read counts.

### MACS2 and SEACR showed different strengths across samples with limited peak overlap

We called peaks using MACS2 and SEACR, the two most widely used peak callers for CUT&RUN data. MACS2 and SEACR are also the default options in many published CUT&RUN pipelines such as CUT&RUNTools [5, 8]. We used each peak caller with its default or recommended parameters to reflect common usage and ensure a fair comparison.

MACS2 performed well across all samples with appropriate processing. SEACR, as intended, worked better on samples with shorter reads and a sparse background, such as the Henikoff Lab samples, but performed poorly on samples with denser background signal.

For some samples, both MACS2 and SEACR performed well. This led us to question whether they identified the same peak sets, as expected if both methods accurately detected the true transcription factor binding sites. To explore this, we counted the number of unique and common peaks between the MACS2 and SEACR peaks in the FOXA2 sample (Figure 4e), considering peaks with at least 50 % overlap as common. Surprisingly, only about 50 % of the peaks overlapped between the two peak callers. Whether this reflects differences in sensitivity, specificity, or underlying peak-calling assumptions remains unclear and requires further investigation.

### SEACR sometimes exhibited large changes in results from minimal changes in fragment length filtering

As noted above, slightly different fragment length cut-offs led to abrupt changes in SEACR’s output (Figure 6). For example, in the C/EBP*β*#1 sample, when using fragments with a length ≤98 bp, SEACR called peaks that showed significant central enrichment of the target motif (concentration: 0.2300). Increasing the cut-off by just 1 bp, however, caused SEACR to call only a single peak, resulting in a concentration value of 0 for the target motif. This sharp discontinuity suggests a lack of robustness in SEACR’s performance under small parameter changes. The FOXA2 sample showed similar behaviour, with concentration at 0.104 for a ≤103 bp cut-off, dropping to 0 when the cut-off decreased by just 1 bp.

**Figure 8.**
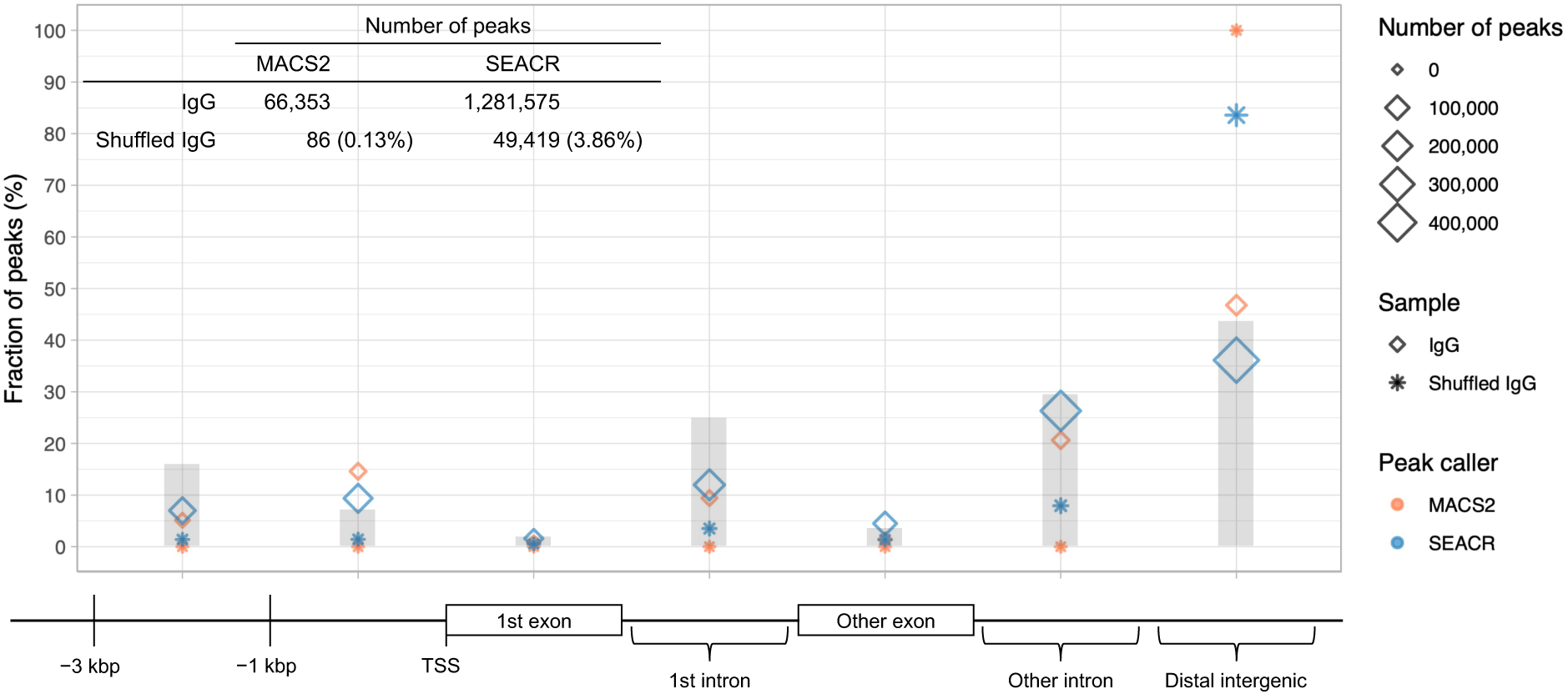
Number of and genomic annotation of peaks called by MACS2 and SEACR on unshuffled and shuffled IgG control data. We randomly shuffled the reads from an IgG control sample and called peaks using the original versus shuffled reads. After shuffling, MACS2 only called 86 peaks while SEACR still called 49 419 peaks. Most of those SEACR peaks mapped to distal intergenic regions, indicating a high level of false positives. Shape: whether the sample was shuffled (star) or not (diamond). Colour hue: peak caller (orange: MACS2, blue: SEACR). Grey bar: fraction of the reference genome covered by each genomic feature. We did not apply fragment length filtering or spike-in calibration for this assessment.

In contrast, MACS2 peak sets showed a steady increase in concentration as the fragment length cut-off decreased, with concentration rising from 0.0468 at 140 bp cut-off to 0.2049 at 90 bp cut-off for the C/EBP*β*#1 sample. Similarly, concentration rose from 0.0611 to 0.1445 for the FOXA2 sample. This trend suggests that MACS2’s performance consistently improved as the fragment length cut-off decreased. This aligns with the expectation that filtering winnowed the remaining fragments down to those containing genuine binding sites.

### MACS2 showed greater robustness than SEACR against calling false-positive peaks from simulated noise

MACS2 and SEACR sometimes produced an unexpectedly large number of peaks, such as more than 100 000 peaks for the C/EBP*β*#2 sample. This raises the question of whether they may identify peaks even when the input data consists entirely of noise. We randomly shuffled the reads from an IgG control sample (Methods). After shuffling, MACS2 called only 86 peaks (0.13 % of the original peak count), within the expected range for false positives. In contrast, SEACR called 49 419 peaks (3.86 % of the original peak count). Most of these peaks mapped to distal intergenic regions (Figure 8), which indicates background noise or false-positive peaks.

We assessed the robustness of both MACS2 and SEACR by increasing the noise level in the input data (Methods). As the noise level increased, MACS2 produced fewer peaks, as expected, ultimately identifying only 691 peaks when the input contained entirely noise (Table 3), which fell within a reasonable range for false positives.

**Table 3.** Mean number of peaks identified by MACS2 or SEACR at six noise levels (percentages) across five simulation replicates. This analysis pertains to the CTCF sample from the Hainer Lab, with noise data generated as described (Methods). MACS2 shows a steady decline in peak numbers with increasing noise. In contrast, SEACR initially calls fewer peaks, but the number increases as the noise level reaches 25%. Bold numbers highlight the largest peak counts observed for each peak caller.

| Noise (%) | MACS2 | SEACR |
| --- | --- | --- |
| 0 | **267 717** | 22 688 |
| 5 | 44 029 | 21 226 |
| 10 | 40 627 | 19 810 |
| 25 | 31 459 | 29 331 |
| 50 | 19 882 | **63 909** |
| 100 | 691 | 54 571 |

In contrast, SEACR calls more peaks as noise levels reach 25 % and continues to identify 54 571 peaks even with only noise (in other words, after shuffling 100 % of reads). This finding aligns with the results from our shuffling experiment, where SEACR similarly calls around 50 000 peaks with entirely shuffled reads. These findings cast doubt on the overall reliability and accuracy of SEACR.

## Discussion

### Overall implications

Overall, we show that applying fragment length filtering with a 120 bp cut-off and proper use of spike-in calibration greatly improves the specificity and centrality of peaks to target motifs. We also assess the robustness of the two popular peak callers for CUT&RUN data, a key consideration when selecting analysis tools. Our work thus serves to better characterize both specific parameters, dataset suitability, and the robustness of each peak caller. While we highlight a number of areas for both further refinement and ongoing improvement, we also provide practical guidance on the optimal use of current analytical methods for this and similar assays.

Careful data processing, including fragment length filtering and spike-in calibration, plays a crucial role in ensuring accurate and reliable interpretation of results. The selection of analysis tools, such as peak callers, should align with specific data characteristics, like background levels, to ensure optimal performance. No single method works best for all datasets; each requires consideration in terms of not only the data in question but also the tool’s overall characteristics and performance.

Our work paints a complex picture of the myriad biochemical and technical considerations and their interplay, which underlie proper analysis of more recent transcription factor binding datasets. We aim not only to provide clear recommendations of practical use to researchers analyzing these datasets, but also to emphasize the underlying biochemical and ensuing statistical complexities that underpin a nuanced analysis of these data. This serves not only to ground their proper interpretation, but can also serve as a foundation upon which to conduct further work, such as the creation of novel algorithms or mathematical models of these distinct assays. To that end, we endeavour to focus here upon various technical considerations, whilst also trying to relate them back to broader implications for this field.

### Study scope and evaluation metric

We focused on transcription factors, where precise identification of binding motifs plays an important role in our understanding of gene regulation. The presence of well-defined sequence motifs also motivated our focus on transcription factors, as motif enrichment can serve as a built-in quality assessment. As such, we did not include chromatin regulators or other DNA-binding proteins that lack known sequence motifs. We likewise excluded datasets targeting histone marks, as these modifications do not associate with discrete DNA motifs and involve broader chromatin features, requiring fundamentally different peak calling strategies. Nonetheless, the approach outlined in this study could inform the later development of algorithms or pipelines suited for histone mark or non-sequence-specific binding data.

To evaluate localized motif enrichment, we chose CentriMo concentration as the primary metric. Concentration measures the total probability assigned to motif occurrences within the defined central region, providing a focused view of motif clustering in narrow genomic windows.^18,19^ This approach aligns with the binding patterns of transcription factors, which often show strong localization to specific regions.^3^ Although CentriMo originated in the context of ChIP-seq, the method itself evaluates positional motif enrichment in equal-length aligned sequences and does not depend on a specific assay.^18^ We therefore used it here as a practical evaluation metric for transcription factor CUT&RUN, where biologically meaningful peaks should likewise show central enrichment of the expected target motif.

E-value remains a widely used metric for assessing the statistical significance of motif enrichment, but prior studies have highlighted challenges with its reliability, particularly in cases of sample size heterogeneity.^33^ These limitations can distort interpretations and rankings based on p-value. E-value rankings, however, still provide valuable insights, particularly when complemented by metrics like concentration that highlight localized motif enrichment. In this study, we compared concentration only across different processing methods within the same sample, avoiding comparisons between samples. Variability in sequencing depth, sample size, and other technical factors makes cross-sample comparisons unreliable. We have investigated this issue more carefully in Viner et al.^33^. While such methods could be applied herein, we find concentration to suffice, especially with such carefully controlled datasets. Still, future work could benefit from the integration of these two, largely orthogonal, approaches.

The p-value ranking in the CentriMo results, which quantitatively ranks motifs enriched at peak centres, serves as another informative metric for motif elucidation.^18^ Ideally, the known target motif would rank first, assuming that biologically relevant motifs tend to show central enrichment near the peak centres. This expectation remains somewhat nuanced, as the assumption may not apply in a number of edge cases and may need to be considered jointly with concentration, as we did here. For instance, many consider a motif’s statistical significance in the CentriMo model quite probative or at least strongly correlated with effect size. As some of us have demonstrated in a related work, however, this does not always hold true—indeed, it can benefit from reinterpretation using more nuanced metrics.^33^ While not the focus of this work, we did try to minimize factors that could complicate interpretation of motif ranks by focusing on concentration and only analyzing highly significant results. We also remained aware of the dangers of over-interpreting small changes in simpler metrics or even their ensuing motif ranks. With respect to fragment-length cut-off selection, one must consider that the target motif rank may remain unchanged despite a slight increase in concentration. The concentration of other similar motifs, such as family motifs, could also increase with fragment length filtering. Under these conditions, a p-value ranking might not provide the most effective way to rank motifs due to heterogeneity in sample size between motifs.^33^

### Fragment length filtering can have complex impacts on binding site elucidation

Filtering for fragments ≤120 bp should enrich for direct transcription factor binding sites, as shorter fragments tend to originate from transcription factor-bound regions rather than nucleosome-protected DNA. This approach aligns with both prior recommendations^1^ and our observations. It may, however, exclude some transcription factor-bound sites that occur in nucleosomal contexts, such as pioneer factor binding sites, because nucleosome-associated DNA cleavage can generate fragments of approximately nucleosomal size. We showed that filtering fragments with a length cut-off of 120 bp could improve motif elucidation, but 120 bp did not serve as an optimal cut-off for all datasets. MACS2 exhibited monotonic improvements as the length cut-off decreased, with fewer called peaks and increased CentriMo concentration of those putatively pertaining to the target motif. Thus, we presume MACS2 exhibited the expected monotonic increase in sensitivity. A data-driven approach to determine the optimal cut-off for each sample helped optimize motif elucidation. Optimizing this cut-off overmuch for MACS2, however, may prove unnecessary, as a slight increase in concentration likely offers minimal benefit.

Returning to the technical issue of response to filtering, we observed several instances where fragment-length filtering reduced SEACR’s performance (Figure 2, NFYB and NFYC). In the filtering experiment for the C/EBP*β*#1 sample, SEACR demonstrated strong performance when applying a fragment length cutoff of 121 bp (Figure 6a). Reducing the cutoff by just 1 bp, however, caused a significant disruption in SEACR’s performance. In both scenarios, the removal of certain fragments sharply increased the threshold SEACR applied for peak calling, which reduced the number of peaks and led to a reduced CentriMo concentration. A possible explanation involves the removal of long fragments, which primarily contribute to background noise under the assumption that short fragments contain true binding sites, while long fragments represent off-target fragments from nucleosome-protected regions. Removing these long fragments could lead to an improved signal-to-noise ratio, due mainly to background signal reduction.

Although counterintuitive at first, the higher threshold applied by SEACR to this cleaner dataset fits its strategy for calibrating thresholds. By filtering the sample to retain shorter fragments, the remaining reads may show stronger signals and minimal noise. This change reshapes the global distribution of signal intensities and amplifies the median signal. In response to this enrichment, SEACR might apply a higher threshold to suppress residual noise and prioritize the most intense peaks. Consequently, the process may remove true peaks with marginally lower signal intensities. The use of a local distribution for peak detection makes MACS2 less sensitive to slight variations in fragment-length cutoff and potentially ameliorates such undesired effects.

### Spike-in calibration factors warrant careful consideration

We identified various factors that influence the results of spike-in calibration. In this study, we use samples with spiked-in DNA fragments from two organisms, *S. cerevisiae* and *E. coli*. Typically, researchers add spike-in DNA to samples before DNA extraction. The organism and preparation of the spike-in DNA fragments could vary widely among laboratories, but they function effectively as long as researchers select spike-in controls carefully, ensuring different DNA sequences and similar G+C contents with the organism of interest.^16^ We demonstrate that calibration using *S. cerevisiae* spike-in DNA can substantially enhance motif elucidation with MACS2. We anticipate that any heterologous spike-in, such as DNA from *Drosophila melanogaster*, would perform similarly well, as long as the spike-in genome remains sufficiently divergent from the target organism to avoid misalignment. We, however, question the use of *E. coli* DNA as a spike-in control due to uncontrolled contamination. For example, MACS2’s performance improved substantially after using both fragment length filtering and spike-in calibration on the NFYB and NFYC samples. The unusually large number of *E. coli* reads in these samples, however, raises concerns about data quality and possible contamination, even if the resulting peak-calling performance appears improved. Similarly, we also have concerns about the validity of using *E. coli* DNA carried over during pA-MNase purification, as proposed in the study that originally introduced this approach as a spike-in control.^25^

We filtered the spike-in reads together with the transcription factor experimental reads for fragment lengths, a step not commonly performed. Filtering the transcription factor reads aligns with the expectation that fragments containing true binding sites typically have lengths of ≤120 bp. On the other hand, we filter the spike-in reads for consistency. The fragment length distribution of spike-in reads varied across samples from different laboratories, suggesting the potential for batch effects or biases in fragment length during library preparation or sequencing. Filtering the spike-in reads clearly improved results for at least one of our samples, and improved or did not harm most others.

As discussed previously, the selected spike-in factor substantially impacts the outcome of MACS2, underscoring the importance of selecting an appropriate coefficient for each sample to ensure accurate normalization. We develop a new spike-in calibration method: instead of using an arbitrary scale, we scale down the larger sample to match the smaller. This approach eliminated the arbitrary nature of the former analysis, which could otherwise cause inconsistencies between samples. This approach represents just one method for performing spike-in calibration, and other methods may offer more accurate calibration between samples. Regardless of the method, spike-in calibration plays a crucial role in ensuring reliable and consistent results across experiments.^16^

The arbitrary scale used in computing spike-in factors greatly influenced MACS2’s performance, but not SEACR’s (Table 2). Since SEACR relies on relative ranking rather than localized noise estimation, it remains stable even when the spike-in factor changes arbitrarily. In particular, ratios between the spike-in factors for the transcription factor and IgG control samples remain constant. This discrepancy arises because MACS2 employs a local background model, dynamically adjusting peak thresholds based on signal-to-background ratios, whereas SEACR applies a global rank-based thresholding approach.

Increasing the scaling factor for spike-in calibration reveals a stepwise pattern in MACS2 performance (Table 2). Using very small scales, like 10, reduces the spike-in factor substantially, leading to a drastic decrease in the normalized signal. With such weak signals, MACS2 calls very few peaks, and the target motif shows no central enrichment. At relatively small scales, such as 50, MACS2 still calls relatively few peaks, and the target motif concentration remains high. This suggests that at low scales, MACS2 can maintain high specificity, due to more stringent peak calling thresholds. As the scaling coefficient increases to values around 100 or 200, MACS2 achieves a balance between sensitivity and specificity. Both the number of peaks and the motif concentration improve, reflecting better detection of genuine target peaks. Our results suggest that this optimal performance occurs because the chosen scales align well with the spike-in read counts in the sample. Once the calibration coefficient surpasses ∼500, however, MACS2 hits a tipping point where the peak count increases dramatically, reaching hundreds of thousands at scales of 5000 or higher, while motif concentration declines substantially. This behaviour results from the inflated calibration coefficient amplifying background noise, leading MACS2 to classify spurious signals as peaks.

This stepwise behaviour in MACS2 likely stems from its use of a local background (noise) model, which dynamically adjusts thresholds based on signal-to-noise ratios. When the calibration coefficient aligns with the spike-in read counts, as seen with scales of ∼100, the model works effectively. Both excessively small scales (such as 10) and excessively large scales (such as 5000), however, disrupt the model, reducing MACS2’s ability to distinguish true peaks from noise. In contrast, SEACR shows no such sensitivity to the arbitrary scale, as it uses a fixed global threshold model. Since the ratio of spike-in factors for the transcription factor and IgG control samples remains constant, SEACR’s performance remained unaffected, regardless of the chosen scaling. These findings emphasize the importance of carefully selecting an appropriate scaling coefficient for spike-in calibration in MACS2 and highlight the utility of our refined empirical scaling factor calibration method.

The calibration strategy also strongly influences the results. Researchers use two main methods for spike-in calibration: one equalizes arbitrary spike-in counts across samples, while the other maintains an equal ratio between spike-in and experimental read counts. We chose the former, as we only performed spike-in calibration between the transcription factor and IgG control samples, not between different biological replicates of the same transcription factor or condition. Substantial differences in the number of experimental read counts between transcription factor samples and IgG controls may occur due to biological reasons. The number of spike-in reads, however, should remain consistent without experimental or sequencing bias, given that the same amount of spike-in DNA goes into each sample.^16^ Hence, ensuring equal counts rather than maintaining an equal ratio between samples better suits our experiments.

We initially hypothesized that differences in mapping specificity between short- and long-read datasets could explain the observed reduction in spike-in reads (Figure 7a). Specifically, short-read datasets may show greater susceptibility to misalignment between the target genome and the spike-in genome. Further examination, however, suggests that mapping specificity alone does not fully account for the observed discrepancies.

Notably, spike-in retention varies across datasets. While De Carvalho and Hainer Lab samples contain comparable spike-in counts before length filtering, the Henikoff Lab samples show a distinct pattern of spike-in depletion, despite having shorter read lengths. This suggests that factors beyond read length, such as differences in spike-in mixing ratios, variability in library preparation steps (such as size selection or ligation efficiency), or sequencing platform-specific biases (such as G+C-content effects or read length preferences), may contribute to the observed differences. This variability highlights an important consideration for spike-in normalization: if recovery rates differ significantly across datasets, normalization may introduce biases rather than reliably correcting for technical variability. Thus, researchers using spike-in normalization for CUT&RUN data should carefully evaluate consistency in spike-in distributions across experimental conditions and consider implementing normalization controls that explicitly account for dataset-specific biases.

### SEACR is highly sensitive to specific dataset parameters

We followed the instructions provided in the manuals for each peak caller to ensure we used the tools as intended. For MACS2, we utilized sub-commands designed for specific tasks, which closely mimic the functionality of the standard `callpeak` command while accommodating spike-in calibration. For SEACR, we used the script with transcription factor and control data, as recommended. While alternative workflows or custom configurations might enhance the performance of these tools, we aimed to evaluate their functionality under standard or suggested conditions, rather than optimizing for the best-case scenario.

We identified a danger zone for CentriMo concentration between 0 and 0.06, where MACS2 calls an excessively large number of peaks (Figure 2, top), many of which include false positives. This issue likely arises due to low-background, zero-dominated signal in CUT&RUN data, which can cause MACS2 to spuriously call weak signals as peaks.^10^ More generally, MACS2 is sensitive to how background is modeled; its original formulation uses a dynamic local background term and can incorporate a control sample to improve specificity. For example, MACS2 achieves higher specificity at the cost of lower sensitivity when using control IgG input data.^34^ In our analyses, fragment length filtering and spike-in calibration prove effective in reducing excessive peaks and improving MACS2’s specificity. SEACR aims to minimize false-positive peak calls and improve specificity in sparse-background data. As shown here, however, SEACR produced a large number of false positives (Figure 2; Table 3). This poses greater challenges for correction through downstream bioinformatic filtering or even through biochemical assay modifications such as increasing read length.

Based upon our observations that small perturbations in fragment length cut-off could dramatically alter SEACR’s results (Figure 6), we initially considered whether an incorrect implementation of its core algorithm might contribute to this unusual behaviour. Further analyses across multiple datasets, however, (both C/EBP*β* samples and FOXA2) suggest that this instability likely does not stem from a software error, but may just reflect an inherent feature of SEACR’s thresholding mechanism. SEACR determines peak significance using a ranked fraction of total signal. Filtering out shorter fragments likely alters the signal distribution in a way that forces a discontinuous threshold shift, rather than a gradual decline in peak detection. Instead of progressively reducing peak calls with increasing fragment length cut-offs, SEACR thus exhibits an abrupt threshold response, dynamically adapting based on the proportion of retained fragments. This behaviour suggests that SEACR’s thresholding approach may exhibit particular sensitivity to signal distribution variability across datasets, leading to inconsistent peak calls when applying strict fragment length filters.

Although SEACR peaks had greater central enrichment for target motifs in several samples compared to MACS2, SEACR’s unpredictable behaviour remains a substantial concern. The lack of robustness and a high false-positive rate make SEACR less reliable for widespread use. We therefore recommend MACS2 as the more reliable option, with proper use of heterologous spike-ins such as *S. cerevisiae*, as it consistently exhibits substantially greater central enrichment across our results.

## Methods

We designed a benchmarking method to evaluate the impact of data processing for CUT&RUN sequencing data, including fragment length filtering, spike-in calibration, and selection of peak callers. We also assessed the robustness of the two peak callers: MACS2^11,12^ and SEACR.^10^

### Datasets

For benchmarking, we curated 12 CUT&RUN datasets to evaluate the performance of peak callers and preprocessing strategies under diverse experimental conditions. To ensure comparability, we selected publicly available datasets from the GEO^20^ database, prioritizing those from different laboratories that met two essential criteria: (1) inclusion of matched IgG control samples, and (2) use of heterologous spike-in DNA for calibration. To assess potential bias, we also incorporated datasets used in the original SEACR publication^10^ (GSM3609741, GSM3609746^25^), generated by the lab that developed both CUT&RUN and SEACR and often cited as assay exemplars. Additionally, we generated new CUT&RUN datasets in K562 cells, targeting C/EBP*β* and ZNF143 with spike-in DNA and matched IgG controls. We have deposited these datasets in GEO (GSE337966).

### Cell culture

We cultured the K562 cells in RPMI-1640 (Hyclone) supplemented with 100 U/mL penicillin (Gibco), 100 µg/mL streptomycin (Gibco), 292 µg/mL L-glutamine (Gibco), and 10 % FBS (Gibco). We maintained the cells at 37 °C and 5 % CO_2_ with routine testing to confirm the lack of Mycoplasma infection.

### CUT&RUN assay

We adapted the CUT&RUN protocol and all buffer recipes from Skene and colleagues^28^ and performed the experiments in 0.2 mL tubes as we recently described.^35^

Briefly, we washed and immobilized 200 000 K562 cells on 10 µL of activated concanavalin A–coated magnetic beads (Polysciences BioMag Plus cat. no. 86057-3). We incubated cells in 0.2 mL antibody buffer containing 1:100 diluted primary antibody. The primary antibodies used in different experiments were anti-C/EBP*β* (Santa Cruz Biotechnology, Dallas, Texas, USA; cat. no. sc-7962), anti-ZNF143 (Proteintech, Rosemont, IL, USA; cat. no. 16618-1-AP; RRID: AB_2218324), or anti-IgG (Abcam, UK; cat. no. ab37415; RRID: AB_2631996). We incubated for 2 h at 4 °C with rotation. Then, we washed the cells with 0.2 mL of digitonin wash buffer. Next, we incubated the cells in 0.2 mL of digitonin wash buffer containing recombinant pA-MNase (expressed and purified in house) diluted to a final concentration of ∼0.8 ng/µL for 1 h at 4 °C with rotation. We then discarded the buffer and replaced it with 150 µL of digitonin wash buffer.

Using an ice-water bath, we cooled the tubes to 0 °C for 5 min and added CaCl_2_ to activate pA-MNase-mediated digestion for 30 min. After 30 min, we added 2× STOP buffer containing chelating agents and heterologous *S. cerevisiae* spike-in DNA to stop the reaction. We incubated the cells at 37 °C for 10 min and centrifuged the cells at 16 000 × g for 5 min at 4 °C. We isolated the DNA from the supernatant containing soluble chromatin fragments via phenol chloroform extraction.

### DNA sequencing

We performed library preparation of 5 ng of CUT&RUN DNA using NEBNext Ultra II DNA Library Preparation Kit for Illumina (NEB, cat. no. E7645L) as per manufacturer’s protocol. We uniquely indexed each sample using NEBNext Multiplex Oligos for Illumina (Index Primers Set 1, cat. no. E7335L; Index Primers Set 2, cat. no. E7500L). We sequenced all libraries at Princess Margaret Genomics Centre (PMGC, Toronto, ON) on an Illumina NextSeq 500 High-Output flow cell using a paired-end 2 × 150 bp read length configuration.

### Overview of CUT&RUN benchmarking method

We used Snakemake^36^ to build a benchmarking pipeline (https://github.com/hoffmangroup/2023cutnrun/) using Python (version 3.7.10). Below, we present a summary of the workflow.

We first trimmed the adapters of paired-end reads and filtered out low-quality reads using the default settings of fastp^37^ (version 0.19.8). Next, we aligned the reads to GRCm38/mm10 or GRCh38/hg38 using Bowtie 2^38^ (version 2.4.4) using options recommended in the CUT&RUN paper.^25^ For samples from the Henikoff Lab, we aligned reads to GRCh37/hg19 to reproduce their work.^10^ While keeping all other parameters the same, we increased the sensitivity of multi-seed alignment by replacing `--very-sensitive` with `-D 20 -R 6 -N 1 -L 18 -i S,1,0.25`. This adjustment increases sensitivity and alignment depth but requires more computational resources.^38^ We selected these parameters to maximize sensitivity while staying within available resources.

The `-N` parameter controls the maximum number of mismatches allowed in the seed region during alignment, directly influencing both specificity and sensitivity.^38^ A setting of `-N=0` enforces strict specificity, allowing only exact seed matches, which reduces ambiguous alignments but may result in lower overall read coverage. Conversely, `-N=1` allows for a single mismatch in the seed region, which increases alignment sensitivity and may improve read retention, particularly in cases of sequencing errors or minor polymorphisms. This adjustment can accordingly enhance the detection of genuine peaks, albeit at the potential cost of increased spurious alignments in repetitive regions, which could affect peak-calling precision.

The paper introducing the CUT&RUN protocol recommended end-to-end mode in Bowtie 2, as CUT&RUN data typically involves short-read sequencing around 25 bp.^25^ This choice ensures a more stringent alignment, minimizing the risk of mismatches that might occur with shorter reads. In our analysis, we applied end-to-end mode for datasets with read lengths ≤80 bp. For longer read lengths (>80 bp), however, we opted for the local mode to better accommodate the increased read length. We also aligned reads to the spike-in genome (sacCer3 for *S. cerevisiae* or MG1655 for *E. coli*) with additional options `--no-overlap --no-dovetail` to avoid possible cross-mapping of the experimental genome to that of the spike-in DNA.

We used Sambamba^39^ (version 0.7.1) for post-processing. We performed fragment length filtering using deepTools alignmentSieve^40^ (version 3.5.1) to retain only reads from fragments with a length ≤120 bp for both the target and spike-in genomes.

We performed spike-in calibration by first counting the spike-in reads in the transcription factor sample and its corresponding IgG control sample. To normalize, we calculated a spike-in factor as the ratio of the smaller spike-in read count to the larger one. We then applied this factor during peak calling to scale down the sample with more spike-in reads, leaving the sample with fewer reads unchanged.

We called peaks with MACS2^11^ (version 2.2.7.1) and SEACR^10^ (version 1.3).

We used the processed BAM files as inputs to MACS2. We used MACS2 subcommands guided by the MACS2 subcommand tutorial (https://github.com/macs3-project/MACS/wiki/Advanced%3A-Call-peaks-using-MACS2-subcommands), with modifications tailored to our analysis. We first generated pileup tracks in BedGraph format using the `pileup` subcommand. To avoid potential errors during the Poisson test, we added a pseudocount of 0.1 using `bdgopt - m add`. We then scaled the signal using the calculated spike-in factor with `bdgopt -m multiply` for spike-in calibration. To estimate local bias from the control sample, we adapted the general strategy described in the tutorial. We created three background tracks using sliding windows of 1 kbp, 10 kbp, and the fragment-length size. Then, we computed the maximum bias for each genomic position by comparing these tracks using `bdgcmp -m max`. When applying spike-in calibration, we included a genome background that represents the average signal of the spike-in scaled transcription factor sample in the maximum bias calculation. Next, we used `bdgcmp -m qpois` to calculate a score track based on the Poisson distribution. Finally, we called peaks using the `bdgpeakcall` subcommand.

We converted the BAM files into BedGraph format required by SEACR using BEDTools^41^ (version 2.27.1) `bamtobed` and `genomecov`. We used R^42^ (version 3.6.1), as required by SEACR. When applying spike-in calibration, we scaled the signal using the calculated spike-in factor with a modified version (https://github.com/hoffmangroup/2023cutnrun/blob/main/scripts/spike_in_calibration_empirical.csh) of the spike-in calibration script from the Henikoff Lab (https://github.com/Henikoff/Cut-and-Run/blob/master/spike_in_calibration.csh). We called SEACR peaks using the stringent mode (by default) without normalization. We skipped the normalization feature in SEACR since we have already performed spike-in calibration, as suggested in the SEACR manual.

We used IGV^43^ and the UCSC Genome Browser^44^ during initial data exploration and visualization. We also used GNU Parallel^45^ throughout our workflow. We used FastQC^46^ (version 0.11.8), Picard^47^ (version 2.10.9) `CollectInsertSizeMetrics`, QualiMap^48^ (version 2.2) `bamqc`, and MultiQC^49^ (version 1.6) for quality control of the sequencing data. We used Java (version 11.0.7), as Picard requires. We used R (version 4.2.1) and ggplot2^50^ (version 3.5.1) for plotting figures.

### Motif analysis and CentriMo plots

We performed motif analyses and created CentriMo graphs using sequences centred on the summit of the peak or the centre of the peak.

For MACS2 peaks, we used the summit position reported in the output narrowPeak files, obtained by adding the `peak` offset to `chromStart`. For SEACR peaks, we used the centre of the maximum-signal region reported in the output BED file as the summit, due to the absence of a clearly defined summit. We used the centre of `chromStart` and `chromEnd` of each peak as the peak centre of both MACS2 peaks and SEACR peaks. For all datasets except those from the Hainer and Henikoff Labs, we used the peak summit as the sequence centre for both MACS2 and SEACR peaks. For datasets from the Hainer and Henikoff Labs, we used the peak summit for MACS2, but the peak centre for SEACR, as the SEACR peak summit did not serve well as a true summit and produced a bimodal motif probability curve.

After determining the centre locus (peak summit or peak centre), we used BEDTools `slop` to extend the region symmetrically to a total length of 500 bp (extending 250 bp upstream and downstream from that locus). We then converted these regions into FASTA format using BEDTools `fastaFromBed`.

We performed motif central enrichment analysis using CentriMo through the MEME-ChIP wrapper command from MEME Suite^51,52^ (version 5.5.2). We used vertebrate motifs from the JASPAR CORE 2020 motif database.^53^ We ran MEME-ChIP with `-meme-nmotifs 0 -streme-nmotifs 0 -fimo-skip -spamo-skip` to restrict the analysis to CentriMo-based central enrichment of known motifs and reduce runtime and memory usage. To avoid overly simple motifs, we used a minimum width of 7, as we found that motifs of length 6 lacked specificity. To avoid motifs with excessive low-information sites, we limited the maximum motif width to 12.

### Spike-in calibration using an arbitrary scale

Unlike the method described above in the benchmarking overview, we additionally performed spike-in calibration using the traditional approach with an arbitrary scale. In the traditional approach, both transcription factor and IgG control samples underwent scaling by multiplying a scale calculated as an arbitrary scale divided by the number of spike-in reads in each sample without length filtering.

### Shuffling experiment

We randomly shuffled reads in the first IgG control sample from the De Carvalho Lab using `randomizeRegions` from R (version 4.2.1) Bioconductor’s^54^ `regioneR`^55^ (version 1.24.0). We called peaks without including any control sample by omitting the `-c` option in MACS2 and applying a numeric threshold of 1 in SEACR. We subsequently conducted motif analyses, as outlined earlier. We annotated peak-containing genomic regions using ChIPseeker^56^ (version 1.34.1) with default settings and the `TxDb.Hsapiens.UCSC.hg38.knownGene` annotation database (version 3.16.0).

We calculated the fraction of the reference genome covered by each genomic feature by summing the widths of all annotated instances of that feature in the annotation database and dividing by the total length of the reference genome.

### Robustness experiment

We randomly shuffled alignment reads in the CTCF sample from the Hainer Lab along the same chromosome using a Python script (https://github.com/hoffmangroup/2023cutnrun/blob/main/scripts/shuffle_entire_genome.py). For each noise level, we combined reads randomly selected from the original sample and the shuffled sample into a single alignment (a merged BAM file) with the script (https://github.com/hoffmangroup/2023cutnrun/blob/main/scripts/shuffle_percentage.py). For example, for 25 % noise, we randomly select 25 % reads from the shuffled sample and 75 % reads from the original sample (Figure 9). Next, we ran our benchmarking workflow to obtain corresponding peak sets from both MACS2 and SEACR. To account for randomness in shuffling and read selection, we repeated the entire process five times and reported the mean number of peaks (Table 3).

**Figure 9.**
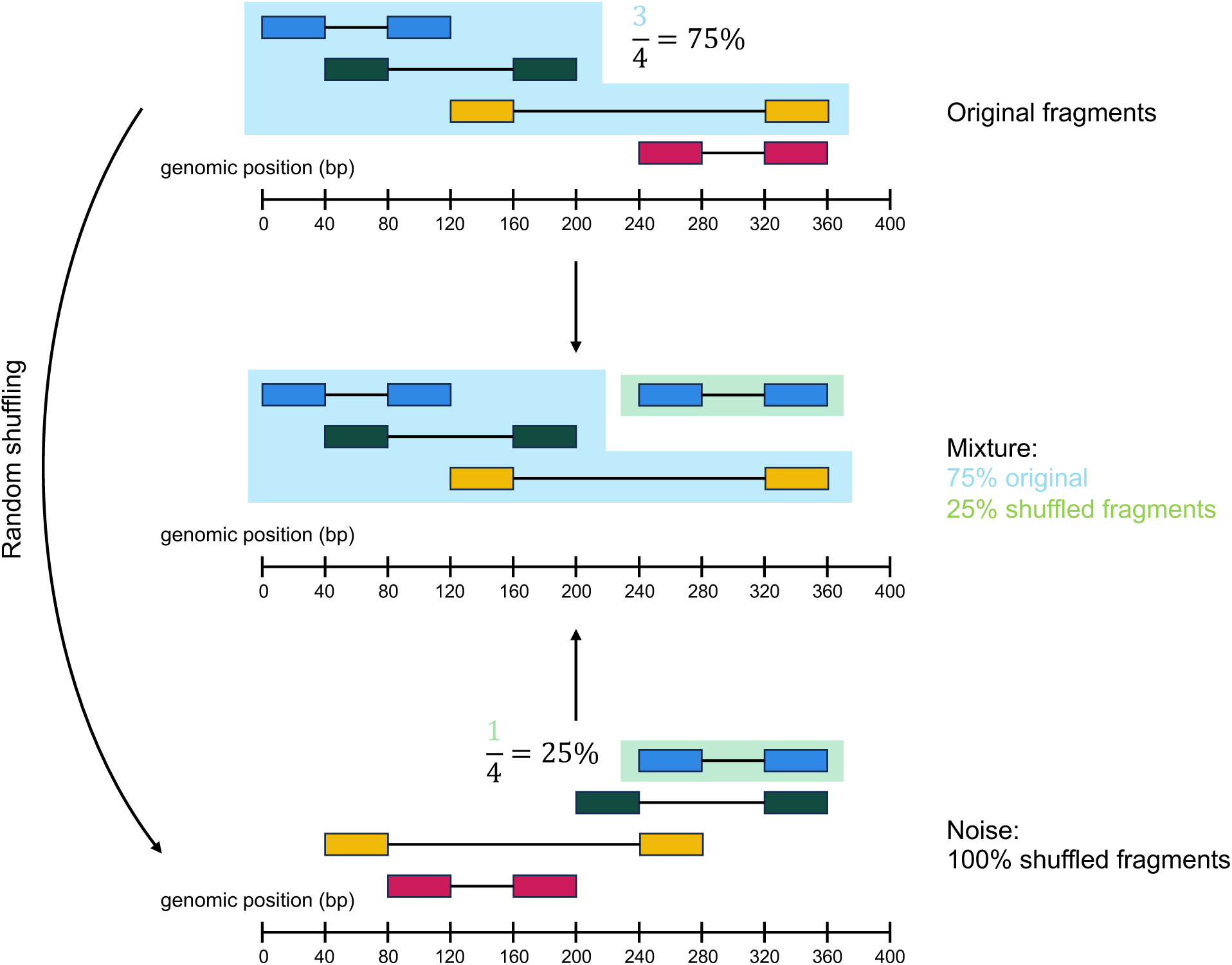
A contrived example to illustrate the generation of noisy data, for our assessment of peak caller robustness. First, each pair of paired-end reads, or fragments, undergoes random shuffling together (on the same chromosome) to generate data with 100% noise. This step changes the positions of the fragments while preserving both fragment lengths and their overall distribution. Next, to generate data with an *x*% noise level, we randomly select a mixture of (100 − *x*)% original fragments and *x*% shuffled fragments (*x* = 25, here). Unlike the fully shuffled case (100% noise), data with partial noise (*x*%) may have a different fragment length distribution compared to the original data due to the sampling mix.

## Software and data availability

The benchmarking pipeline is available on GitHub (https://github.com/hoffmangroup/2023cutnrun/). Persistent availability is ensured by Zenodo, in which we have deposited the version of our code we used (https://doi.org/10.5281/zenodo.21114725) and the peak sets generated for this work (https://doi.org/10.5281/zenodo.21112223). All scripts utilized in this study are publicly available in these repositories and cover all procedures detailed in Methods. All source code is licensed under a [GNU General Public License, version 3 (GPLv3)](https://www.gnu.org/licenses/gpl-3.0.en.html), except for CentriMo, which retains its original license. We have deposited all CUT&RUN sequencing data generated for this work in GEO (GSE337966) We have used a number of published sequencing datasets from GEO: GSM3022415,^23^ GSM3022432,^23^ GSM3609741,^25^ GSM3609746,^25^ GSM2803196,^28^ GSM5703789,^29^ GSM7103770,^30^ GSM7103771,^30^ and GSM7213748.^32^

## Acknowledgments

We thank Timothy L. Bailey for helpful advice regarding the MEME Suite and CentriMo expectations for these datasets. We thank Steven Henikoff (Fred Hutchinson Cancer Center) for the generous sharing of CUT&RUN reagents and experimental guidance. We thank Carl Virtanen and Zhibin Lu (Bioinformatics and High Performance Computing Core, University Health Network) for technical assistance.

## Authors’ contributions

Conceptualization, C.V. and M.M.H.; Data Curation, L.T. and C.V.; Formal Analysis, L.T., C.V., X.H.L., and M.W.; Investigation, L.T., C.V., C.A.I., and S.Y.S.; Methodology, L.T., C.V., X.H.L., M.W., S.J.H., and M.M.H.; Software, L.T., C.V., X.H.L., and M.W.; Visualization, L.T., C.V., and M.M.H.; Writing — Original Draft, L.T. and C.V.; Writing — Review & Editing, L.T., C.V., C.A.I., D.D.De C., S.J.H., and M.M.H.; Resources, D.D.De C. and M.M.H.; Funding Acquisition, M.M.H.; Project Administration, L.T., C.V., and M.M.H.; Supervision, C.V. and M.M.H.

